# Mantis: A Fast, Small, and Exact Large-Scale Sequence-Search Index

**DOI:** 10.1101/217372

**Authors:** Prashant Pandey, Fatemeh Almodaresi, Michael A. Bender, Michael Ferdman, Rob Johnson, Rob Patro

## Abstract

**Motivation:** Sequence-level searches on large collections of RNA-seq experiments, such as the NIH Sequence Read Archive (SRA), would enable one to ask many questions about the expression or variation of a given transcript in a population. Bloom filter-based indexes and variants, such as the Sequence Bloom Tree, have been proposed in the past to solve this problem. However, these approaches suffer from fundamental limitations of the Bloom filter, resulting in slow build and query times, less-than-optimal space usage, and large numbers of false positives.

**Results:** This paper introduces Mantis, a space-efficient data structure that can be used to index thousands of raw-read experiments and facilitate large-scale sequence searches on those experiments. Mantis uses counting quotient filters instead of Bloom filters, enabling rapid index builds and queries, small indexes, and *exact* results, i.e., no false positives or negatives. Furthermore, Mantis is also a colored de Bruijn graph representation, so it supports fast graph traversal and other topological analyses in addition to large-scale sequence-level searches.

In our performance evaluation, index construction with Mantis is 4.4× faster and yields a 20% smaller index than the state-of-the-art split sequence Bloom tree (SSBT). For queries, Mantis is 6× –108× faster than SSBT and has no false positives or false negatives. For example, Mantis was able to search for all 200,400 known human transcripts in an index of 2652 human blood, breast, and brain RNA-seq experiments in one hour and 22 minutes; SBT took close to 4 days and AllSomeSBT took about eight hours.

Mantis is written in `C++11` and is available at https://github.com/splatlab/mantis.

## 1 Introduction

The ability to issue sequence-level searches over publicly available databases of assembled genomes and known proteins has played an instrumental role in many genomics studies, and has made BLAST [2] and its variants some of the most widely-used tools in all of science. Much subsequent work has focused on how to extend tools such as BLAST to be faster, more sensitive, or both [5, 6, 21, 25]. However, the strategies applied by such tools focus on the case where queries are issued over a database of reference sequences. Yet, the vast majority of publicly-available sequencing data (e.g., the data deposited in the SRA [12]) exists in the form of raw, unassembled sequencing reads. As such, this data has mostly been rendered impervious to sequence-level search, which substantially reduces the utility of such publicly available data.

There are a number of reasons that typical reference-database-based search techniques cannot easily be applied in the context of searching raw, unassembled sequences. One major reason is that most current techniques do not scale well as the amount of data grows to the size of the SRA (which today is *≈* 4 petabases of sequence information). A second reason is that searching unassembled sequences means that relatively long queries (e.g., genes) are unlikely to be present in their entirety as an approximate substring of the input.

Recently, new computational schemes have been proposed that hold the potential to allow searching raw sequence read archives while overcoming these challenges. Solomon and Kingsford introduced the sequence Bloom tree (SBT) data structure [22] and an associated algorithm that enables an efficient type of search over thousands of sequencing experiments. Specifically, they re-phrase the query in terms of *k*-mer set membership in a way that is robust to the fact that the target sequences have not been assembled. The resulting problem is coined as the *experiment discovery* problem, where the goal is to return all experiments that contain at least some user-defined fraction *θ* of the *k*-mers present in the query string. The space and query time of the SBT structure has been further improved by Solomon and Kingsford [23] and Sun et al. [26] by applying an all-some set decomposition over the original sets of the SBT structure (Section 2). This seminal work introduced both a formulation of this problem, and the inital steps toward a solution.

Sequence Bloom trees build on prior work using Bloom filters [4]. A Bloom filter is a compact representation of a set *S*. Bloom filters support insertions and membership queries, and they save space by allowing a small false-positive probability. That is, a query for an element *x ∉S* might return “present” with probability *δ*. Allowing false positives enables the Bloom filter to save space—a Bloom filter can represent a set of size *n* with a false-positive probability of *δ* using *O*(*n* log_2_(1*/δ*)) bits. Bloom filters have an interesting property that the bitwise-or of two Bloom filters representing *S*_1_ and *S*_2_ yields a Bloom filter for *S*_1_ *∪ S*_2_. However, the false-positive rate of the union may increase substantially above *δ*.

In *k*-mer-counting tools, Bloom filters are used to filter out single-occurence (and likely erroneous) *k*-mers from raw read data [14]. In a high-coverage genomic data set, any *k*-mer that occurs only once is almost certainly an error and can thus be ignored. However, such *k*-mers can constitute a large fraction of all the *k*-mers in the input—typically 30-50%—so allocating a counter and an entry in a hash table for these *k*-mers can waste a lot of space. Tools such as BFCounter [14] and Jellyfish [13] save space by inserting each *k*-mer into a Bloom filter the first time it is seen. For each *k*-mer in the input, the tool first checks whether the *k*-mer is in the Bloom filter. If not, then this is the first time this *k*-mer has been seen, so it is inserted into the filter. If the *k*-mer is already in the filter, then the counting tool stores the *k*-mer in a standard hash table, along with a count of the number of times this *k*-mer has been seen. In this application, a false positive in the Bloom filter simply means that the tool might count a few *k*-mers that occur only once. Using Bloom filters in this way can reduce space consumption by roughly 50%.

Sequence Bloom trees repurpose Bloom filters to index large sets of raw sequencing data probabilistically. In an SBT, each experiment is represented by a Bloom filter of all the *k*-mers that occur a sufficient number of times in that experiment. A *k*-mer counter can create such a Bloom filter by first counting all the *k*-mers in the experiment and then inserting every *k*-mer that occurs sufficiently often into a Bloom filter. The SBT then builds a binary tree by logically or-ing Bloom filters until it reaches a single root node. To find all the experiments that contain a *k*-mer *x*, start from the root and test whether *x* is in the Bloom filter of each of the root’s children. Whenever a Bloom filter indicates that an element might be present in a subtree, recurse into that subtree.

SBTs support queries for entire transcripts as follows. First compute the set *Q* of *k*-mers that occur in the transcript. Then, when descending down the tree, only descend into subtrees whose root Bloom filter contains at least a *θ* fraction of the *k*-mers in *Q*. Typical values for *θ* proposed by Solomon and Kingsford [22] are in the range 0.7 *−* 0.9. That is, any experiment that contains 70-90% of the *k*-mers in *Q* has a reasonable probability of containing the transcript (or a closely-related variant).

Due to limitations of the Bloom filter, the SBT is forced to balance between the false-positive rate at the root and the size of the filters representing the individual experiments. Because Bloom filters cannot be resized, they must be created with enough space to hold the maximum number of elements that might be inserted. Furthermore, two Bloom filters can only be logically or-ed if they have the same size (and use the same underlying hash functions). Thus, the Bloom filters at the root of the tree must be large enough to represent every *k*-mer in every experiment indexed in the entire tree, while still maintaining a good false positive rate. On the other hand, the Bloom filters at the leaves of the tree represent only a relatively small amount of data and, since there are many leaves, should be as small as possible.

Further complicating the selection of Bloom filter size is that individual dataset sizes vary by orders of magnitude, but all datasets must be summarized using Bloom filters of the same size. For these reasons, most of the Bloom filters in the SBT are, of necessity, sub-optimally tuned and inefficient in their use of space. SBTs partially mitigate this issue by compressing their Bloom filters using an off-the-shelf compressor (Bloom filters that are too large are sparse bit vectors and compress well.) Nonetheless, SBTs are typically constructed with a Bloom filter size that is too small for the largest experiments, and for the root of the tree. As a result, they have low precision: typically only about 57-67% of the results returned by a query are actually valid (i.e., contain a fraction of the query *k*-mers *≥ θ*).

### Results

We present a new *k*-mer-indexing approach, which we call Mantis, that overcomes these obstacles. Mantis has several advantages over prior work:

- Mantis is *exact*. A query for a set *Q* of *k*-mers and threshold *θ* returns exactly those data sets containing at least fraction *θ* of the *k*-mers in *Q*. There are no false positives or false negatives. In contrast, we show that roughly 57-67% of the results in SBT-based systems are false positives.
- Mantis supports much faster queries than existing SBT-based systems. In our experiments, queries in Mantis ran up to 100*×* faster than in SSBT.
- Mantis supports much faster index construction. For example, we were able to build the Mantis index on 2652 data sets in 22 hours. SSBT reported 97 hours to construct an index on the same collection of data sets.
- Mantis uses less storage than SBT-based systems. For example, the Mantis index for the 2652 experiments used in the SSBT evaluation is 20% smaller than the compressed SSBT index for the same data.
- Mantis returns, for each experiment containing at least 1 *k*-mer from the query, the number of query *k*-mers present in this experiment. Thus, the full spectrum of relevant experiments can be analyzed. While these results can be post-processed to filter out those not satisfying a *θ*-query, we believe the Mantis output is more useful, since one can analyze which experiments were close to achieving the *θ* threshold, and can examine if there is a natural “cutoff” at which to filter experiments.

Mantis builds on Squeakr [18], a *k*-mer counter based on the counting quotient filter (CQF) [19]. The CQF is a Bloom-filter alternative that offers several advantages over the Bloom filter. First, the CQF supports counting, i.e., queries to the CQF return not only “present” or “absent,” but also an estimate on the number of times the queried item has been inserted. Analogous to the Bloom filter’s false-positive rate, there is a tunable probability *δ* that the CQF may return a count that is higher than the true count for a queried element. CQFs can also be resized, and CQFs of different sizes can be merged together efficiently. Finally, CQFs can be used in an “exact” mode where they act as a compact exact hash table, i.e., we can make *δ* = 0. CQFs are also faster and more space efficient than Bloom filters.

Prior work has shown how CQFs can be used to improve performance and simplify the design of *k*-mer-counting tools [] and de Bruijn graph representations[]. For example, Squeakr is essentially a thin wrapper around a CQF— it just parses fastq files, extracts the *k*-mers, and inserts them into a CQF. Other *k*-mer counters use multiple data structures (e.g., Bloom filters plus hash tables) and often contain sophisticated domain-specific tricks (e.g., minimizers) to get good performance. Despite its simplicity, Squeakr uses less than half the memory of other *k*-mer counters, offers competitive counting performance, and supports queries for counts up to 10× faster than other *k*-mer counters. Performance is similar in exact mode, in which case, the space is comparable to other *k*-mer counters.

In a similar spirit, Mantis uses the CQF to create a simple space- and time-efficient index for searching for sequences in large collections of experiments. Mantis is based on colored de Bruijn graphs. The “color” associated with each *k*-mer in a colored de Bruijn graph is the set of experiments in which that *k*-mer occurs. We use an exact CQF to store a table mapping each *k*-mer to a color ID, and another table mapping color IDs to the actual set of experiments containing that *k*-mer. Mantis uses an off-the-shelf compressor [20] to store the bit vectors representing each set of experiments.

Mantis takes as input the collection of CQFs representing each data set, and outputs the search index. Construction is efficient because it can use sequential I/O to read the input and write the output CQFs. Similarly, queries for the color of a single *k*-mer are efficient since they require only two table lookups.

We believe that, since Mantis is also a colored de Bruijn graph representation, it may be useful for more than just querying for the existence of sequences in large collections of data sets. Mantis supports the same fast de Bruijn graph traversals as Squeakr, and hence may be useful for topological analyses such computing the length of the query covered in each experiment (rather than just the fraction of *k*-mers present). It can also naturally support operations such as bubble calling [10], and hence could allow a natural, assembly-free way to analyze variation *among* experiments.

## 2 Background and Related Work

The *experiment discovery* problem was first posed by Solomon and Kingsford [22], where they introduced the sequence Bloom tree (SBT). Subsequently, the SBT data structure was improved in two distinct but related works—the Split SBT (SSBT) [23], and AllSomeSBT [26].

The SBT is a binary tree of Bloom filters. Each leaf node is a Bloom filter storing the *k*-mers from a particular experiment. Thus, the number of leaves in the tree equals the number of experiments being indexed. An internal node of the tree is a Bloom filter storing the union of the Bloom filters of its two children. Thus, an internal node’s Bloom filter stores *k*-mers from all the experiments represented in that node’s descendant leaves. In an SBT, all Bloom filters have the same (uncompressed) size. In the published setting [22], each Bloom filter is configured to be 236MB in size, uncompressed. As mentioned earlier (Section 1), the Bloom filters nearer to the root of the SBT have a high error rate, whereas those low in the tree are often nearly empty. This inefficiency is partially mitigated by only inserting a *k*-mer into a leaf if it appears at least a certain number of times in the experiment, so that the total number of indexed *k*-mers is smaller than the total number of distinct *k*-mers. SBTs also support incremental insertion and deletion of individual experiments.

The SSBT and the AllSomeSBT have a similar structure to the SBT, but they use more efficient encodings. The AllSomeSBT has a shorter construction time and query time than the SBT. The SSBT has a slower construction time than the SBT, answers queries faster than the SBT, and uses less space than either the SBT or the AllSomeSBT.

Both structures use a similar high-level approach for saving space and thus making queries fast. Namely, instead of retaining a single Bloom filter at each internal node, the structures maintain two Bloom filters. One Bloom filter stores *k*-mers that appear in every experiment in the descendant leaves. These *k*-mers do not need to be stored in any descendants of the node, thus reducing the space consumption by reducing redundancy. If a queried *k*-mer is found in this Bloom filter, then it is known to be present in all descendant experiments. If the required fraction of *k*-mers for a search (i.e. *θ*) ever appear in such a filter, then search of this subtree can terminate early as all descendant leaf nodes satisfy the query requirements. The other Bloom filter stores the rest of the *k*-mers, those that appear in some, but not all, of the descendants. Both structures save additional space by clustering similar leaves into subtrees, so that more *k*-mers can be stored higher up in the tree and with less duplication.

## 3 Methods

Mantis builds on Squeakr [18], a *k*-mer counter that uses a counting quotient filter [19] as its primary data structure. We first review Squeakr and the CQF, and then we explain how to build upon CQFs to construct our search index.

### 3.1 Squeakr and the counting quotient filter

Squeakr is essentially a thin wrapper around the counting quotient filter. It parses *k*-mers from a collection of input files, and inserts them into a counting quotient filter. It then serializes the CQF to disk.

The CQF is a compact representation of a multi-set *S*, similar in spirit to how a Bloom filter is a compact representation of a set. Thus a CQF supports inserts and queries. A query for an item *x* returns an estimate of the number of instances of *x* in *S*. Like a Bloom filter, a CQF has only one-sided error, i.e., the count returned by a CQF is never smaller than the true count. The CQF also supports a tunable false-positive rate *δ*, which means that a query to the CQF for the count of an item *x* returns the true count of *x* with probability at least 1 *− δ*.

The counting quotient filter represents *S* by storing a compact, lossless representation of the multiset *h*(*S*), where *h*: *U → {*0,…, 2^*p*^ −1*}* is a hash function and *U* is the universe from which *S* is drawn. The CQF sets *p* = log2(*n/δ*) to obtain a false-positive rate *δ* while handling up to *n* insertions [3].

The counting quotient filter saves space by representing some of the bits of *h*(*x*) implicitly using a technique called *quotienting*. The CQF divides *h*(*x*) into its first *q* bits, called the *quotient h*_0_(*x*), and its remaining *r* = *p − q* bits, called the *remainder h*_1_(*x*). It maintains an array *Q* of 2^*q*^ *r*-bit slots, each of which can hold a single remainder. When an element *x* is inserted, the CQF attempts to store the remainder *h*_1_(*x*) in the *home slot Q*[*h*_0_(*x*)]. If that slot is already in use, then the CQF uses a variant of linear probing (using few metadata-bits per slot), to find an unused slot where it can store *h*_1_(*x*) [19].

Instead of storing multiple copies of the same item to count, like a quotient filter, the CQF employs an encoding scheme to count the multiplicity of items. The encoding scheme enables the counting quotient filter to maintain *variable-sized* counters. This is achieved by using slots originally reserved to store the remainders to store count information instead. The metadata bits maintained by the counting quotient filter allows this dynamic reuse of remainder slots for large counters while still ensuring the correctness of all counting quotient filter operations.

The variable-sized counters in the counting quotient filter enable the data structure to handle highly skewed datasets efficiently. By reusing the allocated space, the counting quotient filter avoids wasting extra space on counters and naturally and dynamically adapts to the frequency distribution of the input data. The CQF never takes more space than a quotient filter for storing the same multiset. For highly skewed distributions, like those observed in HTS-based datasets, it occupies only a small fraction of the space that would be required by a comparable (in terms of false-positive rate) quotient filter.

The CQF also supports efficient enumeration of the set *h*(*S*). Enumerating *h*(*S*) involves a linear scan of the quotient filter, so it is both computationally efficient and I/O efficient, if the CQF is stored on disk. This enables efficient merges of several counting quotient filters into into a single filter. We use this functionality during the construction phase of Mantis.

Since the CQF stores *h*(*S*), exactly, the CQF reports inaccurate counts only when there is a collision in *h*. Thus the CQF can be made exact (i.e. *δ* = 0) by using an invertible hash function.

### 3.2 Mantis

Mantis takes as input a collection of experiments and produces a data structure that can be queried with a given *k*-mer to determine the set of experiments containing that *k*-mer. Mantis supports these queries by building a colored de Bruijn graph. In the colored de Bruijn graph, each *k*-mer has an associated color, which is the set of experiments containing that *k*-mer.

The Mantis index is essentially a colored de Bruijn graph, represented using two dynamic data structures: a counting quotient filter and a color-class table. The counting quotient filter is used to map each *k*-mer to a color ID, and then that ID can be looked up in the color-class table to find the actual color (the list of experiments containing that *k*-mer). This approach of using color classes was also used in Rainbowfish [1].

Mantis re-purposes the CQF’s counters to store color IDs instead. In other words, to map a *k*-mer *k* to a color ID *c*, we insert *c* copies of *k* into the CQF. The CQF supports not only insertions and deletions, but directly setting the counter associated with a given *k*-mer, so this can be done efficiently (i.e., we do not need to insert *k* repeatedly to get the counter up to *c*, we can just directly set it to *c*).

Each color class is represented as a bit vector in the color-class table, with one bit for each input experiment (Figure 1). All the bit vectors are concatenated and compressed using RRR compression [20] as implemented in the sdsl library [8].

**Fig. 1:**
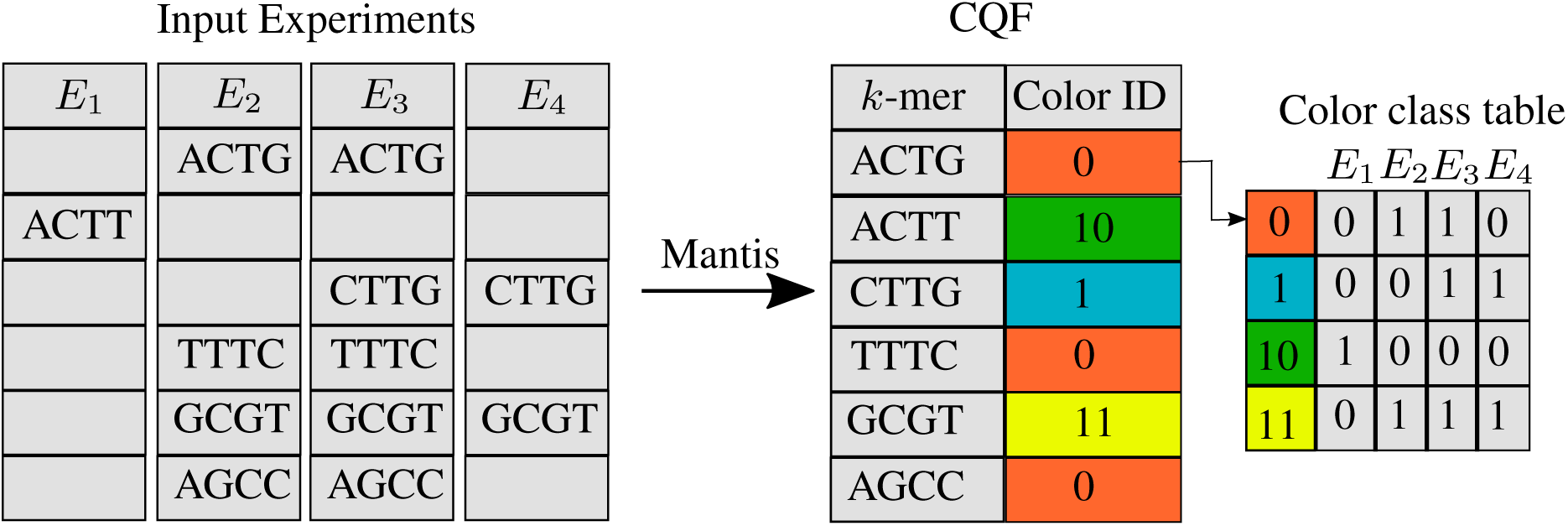
The Mantis indexing data structures. The CQF contains mappings from *k*-mers to color-class IDs. The color-class table contains mappings from color-class IDs to bit vectors. Each bit vector is *N* bits, where *N* is the number of experiments from which *k*-mers are extracted. The CQF is constructed by merging *N* input CQFs each corresponding to an experiment. A query first looks up the *k*-mer(s) in the CQF and then retrieves the corresponding color-class bit vectors from the color-class table.

#### Construction

To construct the Mantis index, we first count *k*-mers for each input experiment using Squeakr. Because Squeakr can either perform exact or approximate *k*-mer counting, the user has the freedom to trade off space and count accuracy. The output of Squeakr is a counting quotient filter containing *k*-mers and their counts. Mantis builds its index by performing a *k*-way merge of the CQFs, creating a single counting quotient filter and a color-class table. The merge process follows a standard *k*-way merge approach [19] with a small tweak.

During a standard CQF merge, the merging algorithm accumulates the total number of occurrences of each key by adding together its counters from all the input CQFs. In Mantis, rather than accumulating the total count of a *k*-mer, we accumulate the set of all input experiments that contain that *k*-mer.

As with SBT-based indexes, once we have computed the set of experiments containing a given *k*-mer, we filter out experiments that contain only a few instances of a given *k*-mer. This filtered set is the color class of the *k*-mer. The merge algorithm then looks up whether it has already seen this color class. If so, it inserts the *k*-mer into the output CQF with the previously assigned color-class ID for this color class. Otherwise, it assigns the next available color-class ID for the new color, adds the *k*-mer’s color class to the set of observed color classes, and inserts the *k*-mer into the output CQF with the new color-class ID. Figure 1 gives an overview of the Mantis build process and indexing data structure.

We keep the uncompressed color-class bit vectors in-memory during the construction process. This helps to quickly look up if a color class has been seen before or not. At the end of the construction process we compress the color-class bit vector using RRR compression [20] and write it to disk.

#### Sampling color classes based on abundance

As explained in Section 3.2, we store the color-class ID corresponding to each *k*-mer as its count in the counting quotient filter. In order to save space in the output CQF, we assign the smaller IDs to the most abundant color classes. We could achieve this using a two-pass algorithm. In the first pass, we count the number of *k*-mers that belong to each color class. In the second pass, we sort the color classes based on their abundances and assign the IDs in increasing order [1]. However, two passes through the data is expensive.

In Mantis, we improve upon the two-pass algorithm as follows. We perform a sampling phase in which we analyze the color-class distribution of a subset of *k*-mers.^3^ We sort the color classes based on their abundances and assign IDs giving the smallest ID to the most abundant color class, an observation that helps in optimizing the size of other data structures like the Bloom Filter Trie [9] and Rainbowfish [1]. We then use this color-class table as the starting point for the rest of the *k*-mers. Given the uniform-randomness property of the hash function used in Squeakr to hash *k*-mers there is a very high chance that we will see the most abundant colors class in the first few million *k*-mers and will assign the smallest ID to it. Figure 2 shows that the smallest ID is assigned to the color class with the most number of *k*-mers. Also, the number of *k*-mers belonging to a color class reduces monotonically as the color class ID increases.

**Fig. 2:**
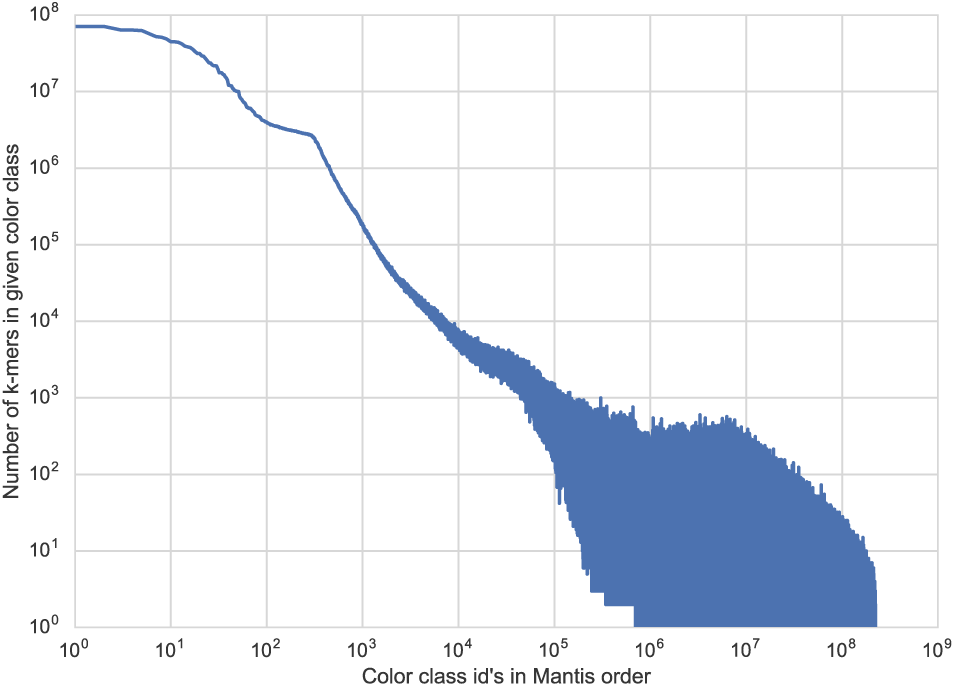
The distribution of the number of *k*-mers in a color class and ID assigned to the color class. The color class with the most number of *k*-mers gets the smallest ID. And the color class with least number of *k*-mers gets the largest ID. This distribution of IDs helps save space in Mantis.

#### Queries

A query consists of a transcript *T* and a threshold *θ*. Let *Q* be the set of the *k*-mers in *T*. The query algorithm should return the set of experiments that contain at least a fraction *θ* of the *k*-mers in *Q*.

Given a query transcript *T* and a threshold *θ*, Mantis first extracts the set *Q* of all *k*-mers from *T*. It then queries the CQF for each *k*-mer *x ∈ Q* to obtain *x*’s color-class ID *c*_*x*_. Mantis then looks up *c*_*x*_ in the color-class table to get the bit vector *v*_*x*_ representing the set of experiments that contain *x*. Finally, Mantis performs vector addition (treating the vectors as vectors of integers) to obtain a single vector *v*, where *v*[*i*] is the number of *k*-mers from *Q* that occur in the *i*th experiment. It then outputs each experiment *i* such that *v*[*i*] *≥ θ|Q|*.

In order to avoid decoding the same color-class bit vector multiple times, we maintain a map from each color-class ID to the number of times a *k*-mer with that ID has appeared in the query. This is done so that, regardless of how many times a *k*-mer belonging to a given color-class appears in the query, we need to decode each color-class at most one time. Subsequently, these IDs are looked up in the color-class table to obtain the bit vector corresponding to the experiments in which that *k*-mer is present. We maintain a hash table that records, for each experiment, the number of query *k*-mers present in this experiment, and the bit vector associated with each color-class is used to update this hash table until the color-classes associated with all query *k*-mers have been processed. This second phase of lookup is particularly efficient, as it scales in the number of *distinct* color-classes that label *k*-mers from the query. For example, if all *n* query *k*-mers belonged to a single color-class, we would decode this color-class’ bit vector only once, and report each experiment present in this color class to contain *n* of the query *k*-mers.

As specified in Section 3.2, Mantis supports both approximate and exact indexes. When used in approximate mode, queries to Mantis may return false positives, i.e. experiments that do not meet the threshold *θ*. Theorem 1 shows that, with high probability, any false positives returned by Mantis are close to the threshold.^4^ Thus, false positives in Mantis are not simply random garbage—they are experiments that contain a significant fraction of the queried *k*-mers.

#### Theorem 1

*A query for q k-mers with threshold θ returns only experiments containing at least θq − O*(*δq* + log *n*) *queried k-mers w.h.p.*

*Proof.* This follows from Chernoff bounds and the fact that the number of queried *k*-mers that are false positives in an experiment is upper bounded by a binomial random variable with mean *δq*.

#### Dynamic updates

We now describe a method that a central authority could use to maintain a large database of searchable experiments. Researchers could upload new experiments and submit queries over previously uploaded experiments. In order to make this service efficient, we need an efficient way to incorporate new experiments without rebuilding the entire index each time a new experiment is uploaded.

Our solution follows the design of cascade filter [19] or LSM-tree [17]. In this approach, the index consists of a logarithmic number of levels, each of which is a single index. The maximum size of each level is a constant factor (typically 4—10) larger than the previous level’s maximum size. New experiments are added to the index at the smallest level (i.e. level 0). Since level 0 is small, it is feasible to add new experiments by simply rebuilding it. When the index at level *i* exceeds its maximum size, we run a merge algorithm to merge level *i* into level *i* + 1, recursively merging if this causes level *i* + 1 to exceed its maximum size.

We now describe how to rebuild a Mantis index (i.e. level 0) to include new experiments. First compute CQFs for the new experiments using Squeakr. Then update the bit vectors in the Mantis index to include a new entry (initially set to 0) for each new experiment. Then run a merge algorithm on the old index and the CQFs of the new experiments. The merge will take as input the old Mantis CQF, the old Mantis mapping from color IDs to bit vectors, and the new CQFs, and will produce as output a new CQF and new mapping from color IDs to bit vectors.

For each *k*-mer during the merge, compute the new set of experiments containing that *k*-mer by adding any new experiments containing that *k*-mer to the old bit vector for that *k*-mer. Then assign that set a color ID as described above and insert the *k*-mer and its color ID into the output CQF.

Merging two levels together is similar. During the merge, simply union the sets of experiments containing a given *k*-mer. This process is I/O efficient since it requires only sequentially reading each of the input indexes.

During a query, we have to check each level for the queried *k*-mers. However, since queries are fast and there are only a logarithmic number of levels, performance should still be good (and, according to our experiments, still better than other state-of-the-art sequence-level search data structures).

## 4 Evaluation

Our evaluation aims to compare Mantis to the state of the art, SSBT [23], on the following performance metrics:

- **Construction time**. How long does it take to build the index?
- **Index size**. How large is the index, in terms of storage space?
- **Query performance**. How long does it take to execute queries?
- **Quality of results**. How many false positives are included in query results?

### 4.1 Experimental procedure

For all experiments in this paper, unless otherwise noted, we consider the *k*-mer size to be 20 to match the parameters adopted by Solomon and Kingsford [22].

To facilitate comparison with SSBT, we use the same set of 2652 experiments used in the evaluation done by Solomon and Kingsford [24], and as listed on their website [11]. These experiments consist of RNA-seq short-read sequencing runs of human blood, brain, and breast tissue. We obtained these files directly from NIH SRA [16]. We discarded 66 files that contained only extremely short reads (i.e., less than 20 bases).^5^ Thus the actual number of files used in our evaluation was 2586.

We first used Squeakr-exact to construct counting quotient filters for each experiment. We used 40-bit hashes in Squeakr-exact to represent *k*-mers exactly. Before running Squeakr-exact, we needed to select the size of the counting quotient filter for each experiment. We used the following rule of thumb to estimate the counting quotient filter size needed by each experiment: singleton *k*-mers take up 1 slot in the counting quotient filter, doubletons take up 2 slots, and almost all other *k*-mers take up 3 slots. We implemented this rule of thumb as follows. We used ntCard [15] to estimate the number of distinct *k*-mers *F*_0_ and the number of *k*-mers of count 1 and 2 (*f*_1_ and *f*_2_, respectively) in each experiment. We then estimated the number of slots needed in the counting quotient filter as *s* = *f*_1_ + 2*f*_2_ + 3(*F*_0_ *− f*_1_ − *f*_2_). The number of slots in the counting quotient filter must be a power of 2, so let *s*′ be the smallest power of 2 larger than or equal to *s*. In order to be robust to errors in our estimate, if *s* was more than 0.8*s*′, then we constructed the counting quotient filter with 2*s*′ slots. Otherwise, we constructed the counting quotient filter with *s*′ slots.

We then used Squeakr-exact to construct a counting quotient filter of the counts of the *k*-mers in each experiment. The total size of all the counting quotient filters was 2.7TB.

We then computed cutoffs for each experiment according to the rules defined in the SBT paper [23], shown in Table 1. The SBT paper specifies cutoffs based on the size of the experiment, measured in bytes, but does not specify how the size of an experiment is calculated (e.g. compressed or uncompressed). We use the size of the compressed file downloaded from SRA.

**Table 1:** Minimum number of times a *k*-mer must appear in an experiment in order to be counted as abundantly represented in that experiment (taken from the SBT paper).

| Min size | Max size | Cutoff |
| --- | --- | --- |
| 0 | $\leq 300\text{MB}$ | 1 |
| $> 300\text{MB}$ | $\leq 500\text{MB}$ | 3 |
| $> 500\text{MB}$ | $\leq 1\text{GB}$ | 10 |
| $> 1\text{GB}$ | $\leq 3\text{GB}$ | 20 |
| $> 3\text{GB}$ | ∞ | 50 |

We then invoked Mantis to build the search index using these cutoffs. Based on a few trial runs, we estimated that the number of slots needed in the final counting quotient filter would be 2^34^, which turned out to be correct.

#### Query datasets

For measuring query performance, we randomly selected sets of 10, 100, and 1000 transcripts from the Gencode annotation of the human transcriptome [7]. Also, before using these transcripts for queries we replaced any occurrence of “N” in the transcripts with a pseudo-randomly chosen valid nucleotide. We then performed queries for these three sets of transcripts in both Mantis and SSBT. For SSBT, we used *θ* = 0.7, 0.8, and 0.9. For Mantis, *θ* makes no difference to the run-time, since it is only used to filter the list of experiments at the very end of the query algorithm. Thus performance was indistinguishable, so we only report one number for Mantis’s query performance.

For SSBT, we used the tree provided to us by the SSBT authors via personal communication.

We also compared the quality of results from Mantis and SSBT. Mantis is an exact representation of *k*-mers and therefore all the experiments reported by Mantis should also be present in the results reported by SSBT. However, SSBT results may contain false-positive experiments. Therefore, we can use Mantis to empirically calculate the precision of SSBT. Precision is defined as *TP/*(*TP* + *FP*), where *TP* and *FP* are the number of true and false positives, respectively, in the query result.

### 4.2 Experimental setup

All experiments were performed on an Intel(R) Xeon(R) CPU (E5-2699 v4 @2.20GHz with 44 cores and 56MB L3 cache) with 512GB RAM and a 4TB TOSHIBA MG03ACA4 ATA HDD running ubuntu 16.10, and were carried out using a single thread. The data input to the construction process (i.e., fastq files and the Squeakr representations) was stored on 4-disk mirrors (8 disks total), each is a Seagate 7200rpm 8TB disk (ST8000VN0022). They were formatted using ZFS and exported to via NFS over a 10Gb link.

The time reported for construction and query benchmarks is the total time taken measured as the wall-clock time using `/usr/bin/time`.

We compare Mantis and SSBT on their in-memory query performance. For Mantis, we warmed the cache by running the query benchmarks twice; we report the numbers from the second run. We followed the SSBT author’s procedure for measuring SSBT’s in-RAM performance [24], as explained to us in personal communication. We copied all the nodes in the tree to a ramfs (i.e. an in-RAM file system). We then ran SSBT on the files stored in the ramfs. (We also tried running SSBT twice, as with Mantis, and performance was identical to that from following the SSBT authors’ procedures.)

SSBT query benchmarks were run with thresholds 0.7, 0.8 and 0.9, and the `max-filter` parameter was set to 11000. By setting `max-filter` to 11000, we ensured that SSBT never had to evict a filter from its cache.

All experiments were performed on an Intel(R) Xeon(R) CPU (E5-2699 v4 @2.20GHz with 44 cores and 56MB L3 cache) with 512GB RAM and a 4TB TOSHIBA MG03ACA4 ATA HDD running ubuntu 16.10, and were carried out using a single thread. The data input to the construction process (i.e., fastq files and the Squeakr representations) was stored on 4-disk mirrors (8 disks total), each is a Seagate 7200rpm 8TB disk (ST8000VN0022). They were formatted using ZFS and exported to via NFS over a 10Gb link.

### 4.3 Results

In this section we present our benchmark results comparing Mantis and SSBT to answer questions posed in Section 4.

#### Build time and index space

Table 2 shows that Mantis builds its index 4.4*×* faster than SSBT. Also, the space needed by Mantis to represent the final index is 20% smaller than SSBT.

**Table 2:**
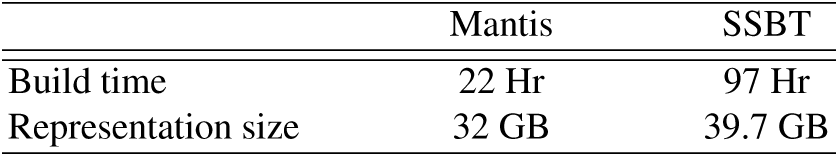
Time and space measurement for Mantis and SSBT. Total time taken by Mantis and SSBT to construct the representation. Total space needed to store the representation by Mantis and SSBT. Numbers for SSBT were taken from the SSBT paper [23].

In Mantis the time spent in compression phase in which we compress the color-class bit vectors is only a small fraction of the total time. Most of the time is spent in merging the hashes from the input counting quotient filters and creating color-class bit vectors. The merging process is fast because there is no random disk IO. We read through the input counting quotient filters sequentially and also insert hashes in the final counting quotient filter sequentially. The number of *k*-mers in the final counting quotient filter was *≈* 3.69 billion.

#### Query performance

Table 3 shows the query performance of Mantis and SSBT on three different query datasets. Even for *θ* = 0.9 (the best case of SSBT), Mantis is 5×, 16×, and 75× faster than SSBT for in-memory queries. For *θ* = 0.7, Mantis is up to 137 times faster than SSBT.

**Table 3:**
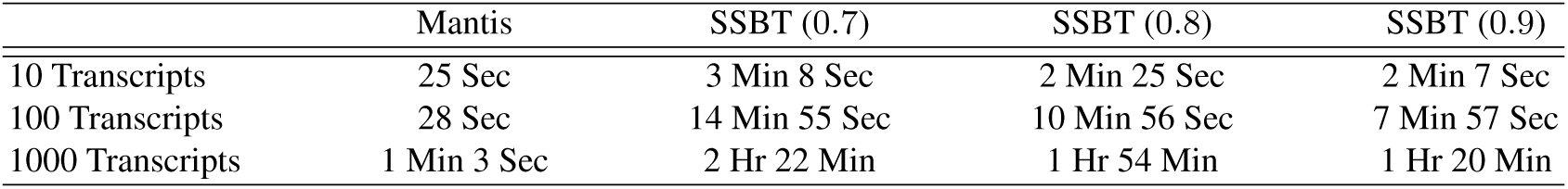
Time taken by Mantis and SSBT to perform queries on three sets of transcripts. The set sizes are 10, 100, and 1000 transcripts. For SSBT we used three different threshold values 0.7, 0.8, and 0.9. All the experiments were performed by either making sure that the index structure is cached in RAM or is read from ramfs.

The query time for SSBT reduces with increasing *θ* values because with higher *θ* queries will terminate early and have to perform fewer accesses down the tree. In Table 3, for Mantis we only have one column because Mantis reports experiments for all *θ* values.

In Mantis, only two memory accesses are required per *k*-mer—one in the counting quotient filter and, if the *k*-mer is present, then the ID is looked up in the color-class table. SSBT has a fast case for queries that occur in every node in a subtree or in no node of a subtree, so it tends to terminate quickly for queries that occur almost everywhere or almost nowhere. However, for random transcript queries, it may have to traverse multiple root-to-leaf paths, incurring multiple memory accesses. This can cause multiple cache misses, resulting in slow queries.

#### Quality of results

Table 4 shows how the results of the queries performed in Section 4.3 compare for Mantis and SSBT. All the comparisons were performed for *θ* = 0.8. As explained in Section 4.1, we use the results from Mantis to calculate the precision in SSBT results. The precision in SSBT result varies from 0.577 to 0.679.

**Table 4:**
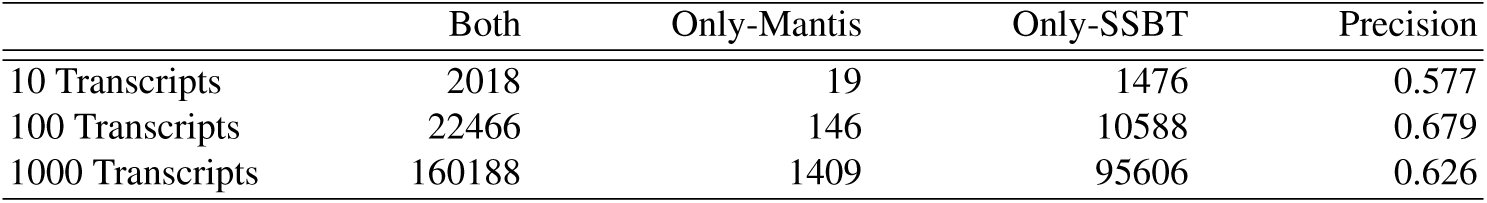
Comparison of query benchmark results for Mantis and SSBT. Both means the number of those experiments that are reported by both Mantis and SSBT. Only-Mantis and Only-SSBT means the number of experiments reported by only Mantis and only SSBT. All three query benchmarks are taken from Table 3 for *θ* = 0.8.

Mantis is an exact index and SSBT is an approximate index with one-sided error (i.e., only false positives). Therefore the experiments reported by SSBT should be a super-set of the experiments reported by Mantis. However, when we compare the results from Mantis and SSBT there are a few experiments reported by Mantis that are not reported by SSBT. These could be because of a bug in Mantis or SSBT.

In order to determine whether there was a bug in Mantis, we randomly picked a subset of experiments reported only by Mantis and used KMC2 to validate Mantis’ results. The results reported by KMC2 were exactly the same as the results reported by Mantis. This means that there were some experiments that actually had at least 80% of the *k*-mers from the query but were not reported by SSBT.

We contacted the SSBT authors about this anomaly, and they found that some of the data sets were corrupted during download from SRA. This resulted in the SSBT-tree having corrupted data for those particualar data sets, which was revealed by comparison with results from Mantis. We believe only a handful of data sets were corrupted, so that these issues do not materially impact the results reported in the SSBT paper.

## 5 Discussion and Conclusion

We have introduced Mantis, a new method and system for tackling the *experiment discovery* (i.e. large-scale sequence search) problem. Though inspired by the Sequence Bloom Tree and subsequent work, Mantis takes a completely different approach to this problem. Specifically, rather than adopting a hierarchy of Bloom filters, as suggested by previous approaches[22, 23, 26], we build our system on top of the counting quotient filter [19]; using this data structure both for counting and as a general key-value store. We combine this data structure with a color-encoding scheme similar to that adopted by Almodaresi et al. for colored de Bruijn graph representation [1]. This different approach allows Mantis to represent, in similar memory to the Split Sequence Bloom Tree [23] (SSBT), a data structure for rapid and *exact k*-mer search over thousands of experiments. Specifically, we have shown that Mantis can be constructed efficiently, that the final structure takes slightly less space than the compressed SSBT, and that it can be queried very rapidly. Moreover, since the representation of Mantis is exact, it exhibits no false positive results. Even though the false positive *rate* of SBT-based solutions is generally low (false positives occur at a typical rate of *∼* 5% in our experiments), this can still translate into a considerable number of experiments when the query space is large. Thus, Mantis represents an attractive methodology for indexing experiments and for addressing the experiment discovery problem. For a similar space requirement as the best-in-class solution, it is faster to construct, provides considerably faster queries, and is exact where other systems are approximate.

We note that Mantis can easily be made approximate, and that this can be done with a tightly-controlled error rate (Theorem 1). In the future, it will be interesting to scale Mantis to even larger collections of data, and to experiment with approximate versions of the system. Also, given that Mantis encodes what is essentially a colored de Bruijn graph over all of the indexed experiments, future work will explore other uses of this representation apart from experiment discovery. For example, Mantis could be used to efficiently search for and categorize variants among unassembled experiments by adopting the “bubble-calling” algorithm over colored de Bruijn graphs proposed by Iqbal et al. [10]. Finally, it could be promising to combine certain concepts from Mantis with ideas from SBT-based solutions—specifically the idea of making the indexing structure hierarchical. Good partitionings of the experimental data, if they can be discovered efficiently, could lead to an even more compact index.

## 6 Acknowledgments

We gratefully acknowledge support from NSF grants BBSRC-NSF/BIO-1564917, IIS-1247726, IIS-1251137, CNS-1408695, CCF-1439084, CCF-1617618, CCF-1716252, and from Sandia National Laboratories.

## Bibliography

[1] Fatemeh Almodaresi, Prashant Pandey, and Rob Patro. Rainbowfish: A Succinct Colored de Bruijn Graph Representation. In Russell Schwartz and Knut Reinert, editors, 17th International Workshop on Algorithms in Bioinformatics (WABI 2017), volume 88 of Leibniz International Proceedings in Informatics (LIPIcs), pages 18:1–18:15, Dagstuhl, Germany, 2017. Schloss Dagstuhl–Leibniz-Zentrum fuer Informatik.

[2] Stephen F Altschul, Warren Gish, Webb Miller, Eugene W Myers, and David J Lipman. Basic local alignment search tool. Journal of molecular biology, 215(3):403–410, 1990.

[3] Michael A. Bender, Martin Farach-Colton, Rob Johnson, Russell Kaner, Bradley C. Kuszmaul, Dzejla Medjedovic, Pablo Montes, Pradeep Shetty, Richard P. Spillane, and Erez Zadok. Don’t thrash: How to cache your hash on flash. Proceedings of the VLDB Endowment, 5(11), 2012.

[4] Burton H. Bloom. Space/time trade-offs in hash coding with allowable errors. Commun. ACM, 13(7):422–426, 1970.

[5] Benjamin Buchfink, Chao Xie, and Daniel H Huson. Fast and sensitive protein alignment using DIAMOND. Nature methods, 12(1):59–60, 2015.

[6] Noah M Daniels, Andrew Gallant, Jian Peng, Lenore J Cowen, Michael Baym, and Bonnie Berger. Compressive genomics for protein databases. Bioinformatics, 29(13):i283–i290, 2013.

[7] Gencode. Release 25. https://www.gencodegenes.org/releases/25.html, 2017. [online; accessed 06-Nov-2017].

[8] Simon Gog. Succinct data structure library. https://github.com/simongog/sdsl-lite, 2017. [online; accessed 01-Feb-2017].

[9] Guillaume Holley, Roland Wittler, and Jens Stoye. Bloom filter trie: an alignment-free and reference-free data structure for pan-genome storage. Algorithms Mol. Biol., 11:3, 2016.

[10] Iqbal Zamin, Caccamo Mario, Turner Isaac, Flicek Paul, and McVean Gil. De novo assembly and genotyping of variants using colored de Bruijn graphs. Nature Genetics, 44:226–232, January 2012.

[11] Carl Kingsford. Srr list. https://www.cs.cmu.edu/~ckingsf/software/bloomtree/srr-list.txt, 2017. [online; accessed 06-Nov-2017].

[12] Yuichi Kodama, Martin Shumway, and Rasko Leinonen. The sequence read archive: explosive growth of sequencing data. Nucleic acids research, 40(D1):D54–D56, 2011.

[13] Guillaume Marçais and Carl Kingsford. A fast, lock-free approach for efficient parallel counting of occurrences of k-mers. Bioinformatics, 27(6):764–770, 2011.

[14] Pall Melsted and Jonathan K Pritchard. Efficient counting of k-mers in dna sequences using a Bloom filter. BMC bioinformatics, 12(1):1, 2011.

[15] Hamid Mohamadi, Hamza Khan, and Inanç Birol. ntcard: a streaming algorithm for cardinality estimation in genomics data. Bioinformatics, 33(9):1324–1330, 2017.

[16] NIH. Sra. https://www.ebi.ac.uk/ena/browse, 2017. [online; accessed 06-Nov-2017].

[17] Patrick O’Neil, Edward Cheng, Dieter Gawlic, and Elizabeth O’Neil. The log-structured merge-tree (LSM-tree). Acta Informatica, 33(4):351–385, 1996.

[18] Prashant Pandey, Michael A Bender, Rob Johnson, and Rob Patro. Squeakr: An exact and approximate k-mer counting system. Bioinformatics, page btx636, 2017.

[19] Prashant Pandey, Michael A. Bender, Rob Johnson, and Robert Patro. A general-purpose counting filter: Making every bit count. In Semih Salihoglu, Wenchao Zhou, Rada Chirkova, Jun Yang, and Dan Suciu, editors, Proceedings of the 2017 ACM International Conference on Management of Data, SIGMOD Conference 2017, Chicago, IL, USA, May 14-19, 2017, pages 775–787. ACM, 2017.

[20] Rajeev Raman, Venkatesh Raman, and S. Srinivasa Rao. Succinct indexable dictionaries with applications to encoding k-ary trees and multisets. In David Eppstein, editor, Proceedings of the Thirteenth Annual ACM-SIAM Symposium on Discrete Algorithms, January 6-8, 2002, San Francisco, CA, USA., pages 233–242. ACM/SIAM, 2002.

[21] Michael Remmert, Andreas Biegert, Andreas Hauser, and Johannes Söding. Hhblits: lightning-fast iterative protein sequence searching by hmm-hmm alignment. Nature methods, 9(2):173–175, 2012.

[22] Brad Solomon and Carl Kingsford. Fast search of thousands of short-read sequencing experiments. Nature biotechnology, 34(3):300–302, 2016.

[23] Brad Solomon and Carl Kingsford. Improved search of large transcriptomic sequencing databases using split sequence bloom trees. In International Conference on Research in Computational Molecular Biology, pages 257–271. Springer, 2017.

[24] Brad Solomon and Carl Kingsford. Improved search of large transcriptomic sequencing databases using split sequence bloom trees. In Süleyman Cenk Sahinalp, editor, Research in Computational Molecular Biology - 21st Annual International Conference, RECOMB 2017, Hong Kong, China, May 3-7, 2017, Proceedings, volume 10229 of Lecture Notes in Computer Science, pages 257–271, 2017.

[25] Martin Steinegger and Johannes Söding. MMseqs2 enables sensitive protein sequence searching for the analysis of massive data sets. Nature biotechnology, 2017.

[26] Chen Sun, Robert S Harris, Rayan Chikhi, and Paul Medvedev. Allsome sequence bloom trees. In International Conference on Research in Computational Molecular Biology, pages 272–286. Springer, 2017.

